# Amphiphilic particle-stabilized nanoliter droplet reactors with a multi-modal portable reader for distributive biomarker quantification

**DOI:** 10.1101/2023.04.24.538181

**Authors:** Vishwesh Shah, Xilin Yang, Alyssa Arnheim, Shreya Udani, Derek Tseng, Yi Luo, Mengxing Ouyang, Ghulam Destgeer, Omai Garner, Hatice Koydemir, Aydogan Ozcan, Dino Di Carlo

## Abstract

Compartmentalization, leveraging microfluidics, enables highly sensitive assays; but the requirement for significant infrastructure for their design, build, and operation limits access. Newer multi-material particle-based technologies thermodynamically stabilize monodisperse droplets as individual reaction compartments with simple liquid handling steps, precluding the need for expensive microfluidic equipment. Here, we further improve the accessibility of this lab on a particle technology to resource-limited settings by combining this assay system with a portable multi-modal reader, thus enabling nanoliter droplet assays in an accessible platform. We show the utility of this platform in measuring N-terminal propeptide B-type natriuretic peptide (NT-proBNP), a heart failure biomarker, in complex medium and patient samples. We report a limit of detection of ∼0.05 ng/ml and a linear response between 0.2 – 2 ng/ml in spiked plasma samples. We also show that, owing to the plurality of measurements per sample, “swarm” sensing acquires better statistical quantitation with a portable reader. Monte Carlo simulations show the increasing capability of this platform to differentiate between negative and positive samples, i.e. below or above the clinical cut-off for acute heart failure (∼0.1ng/ml), as a function of the number of particles measured. Our platform measurements correlate with gold standard ELISA measurement in cardiac patient samples, and achieve lower variation in measurement across samples compared to the standard well plate-based ELISA. Thus, we show the capabilities of a cost-effective droplet-reader system in accurately measuring biomarkers in nanoliter droplets for diseases that disproportionally affect underserved communities in resource-limited settings.

---

The ability to compartmentalize reactions into extremely small volumes has enabled the study of biology with exquisite sensitivity (Basu, 2017a, 2017b). Low/no crosstalk and uniformity ensure each partition serves as an individual reaction compartment with comparable conditions. Currently available compartmentalization technologies, based on droplets or microwell arrays, require specialized infrastructure to operate (Rissin et al., 2010; Rondelez et al., 2005; Yelleswarapu et al., 2019). Droplet assays are also marred by surfactant-induced diffusion of reaction products leading to crosstalk (Gruner et al., 2016). These limit their accessibility and usability for diagnostics and research.

Previously, we described the production and use of multi-material amphiphilic particles made of concentric hydrophobic and hydrophilic polymer layers to thermodynamically stabilize water-in-oil emulsions.(Destgeer et al., 2020; Wu et al., 2020) This lab on a particle method leverages the surface interactions of two different polymers to create a local energy minimum in the volume-energy curves, thereby stabilizing a fixed volume of the aqueous solution. This method of generating monodisperse reaction volumes with simple mixing steps democratizes sensitive droplet assays for use in life sciences research and clinical diagnostics. Previous work, utilizing these particles with readout on benchtop inverted microscopes, reported limits of detection of NT-proBNP, a canonical heart failure marker, down to femtogram per milliliter concentrations in buffer testing (Destgeer et al., 2020).

The need for benchtop fluorescence and brightfield microscopes for end-point readout limits the deployment of this otherwise accessible technology in local clinics and resource-limited settings. Not surprisingly, the populations in these low-resourced regions are often disproportionately affected by otherwise preventable medical conditions, and could benefit from technologies for sensitive and quantitative biomarker detection. For example, historical and ongoing evidence shows that heart failure (HF) disproportionately negatively affects underserved populations in economically deprived regions often comprising of minority populations (Adam Leigh et al., 2016; Benjamin et al., 2019; Mozaffarian et al., 2015; Virani et al., 2021). The Multi-Ethnic Study of Atherosclerosis found that African American and Hispanic populations in the United States had the highest risk of developing HF, and African Americans had the highest proportion of incident HF not preceded by myocardial infarction (75%) (Bahrami et al., 2008). While HF is treatable upon diagnosis, a recent study found that ∼80% of patients are only diagnosed upon emergency hospital admission (Taylor et al., 2021). AHA guidelines establish the measurement of N-terminal propeptide B-type natriuretic peptide (NT-proBNP) for the diagnosis of HF with a clinical cutoff of 0.125 ng/ml. (Yancy et al., 2017) Results from the Heart Failure Assessment With BNP in the Home (HABIT) showed that B-type Natriuretic Peptide (BNP) levels, monitored as a continuous variable over time, can be correlated to the risk of a patient experiencing acute clinical heart failure decompensation (ADHF) (Maisel et al., 2013); thus can help reduce 30-day emergency hospital readmission rates. Improving access to NT-proBNP testing at local clinics and other outpatient settings can help detect HF before emergency hospital admission and improve patient outcomes, especially in underserved communities.

Here, we present a cost-effective lab on a particle-based assay and develop a portable reader and algorithm to measure NT-proBNP levels in plasma-EDTA and patient samples. The microparticle-based assay achieves amplified sensing in tens to hundreds of individual compartmentalized reaction volumes simultaneously, resulting in enhanced performance through increased statistical sampling, and follows a simple workflow. The portable, low-cost reader has a small footprint (10” x 8”) and can be placed in local clinics and remote health centers. The images generated for an entire reaction well are analyzed by the portable reader to detect each particle and the reaction occurring within the enclosed droplets. Leveraging this platform, we show ratiometric response to increasing doses of NT-proBNP spiked into plasma-EDTA sample, and a correlated measurement between this platform and standard well plate ELISA in cardiac patient samples. We conducted Monte Carlo simulations showing the decreasing error rates in measurements, and increasing capability to distinguish between samples at and above the clinical cut-off for acute heart failure (0.1ng/ml), and those below as a function of the number of simultaneous particle measurements obtained from each sample.

## Results and Discussion

### Amphiphilic Lab-on-a-Particle ELISA

The assay uses multi-material particles (∼300 µm in diameter) with concentric layers of an inner polyethylene glycol (PEG) layer and an outer polypropylene glycol (PPG) layer with a cavity in the middle (fig. 1a), which allows them to hold a small aqueous reaction volume. Particles were fabricated using 3D-printed microfluidic devices that are smaller and redesigned structurally (supplementary fig. 1) for more streamlined fabrication compared to those previously described, possible by advances in micro stereolithography (Destgeer et al., 2020, 2021). During fabrication, the inner PEG region was functionalized with biotin to enable linking to streptavidin-conjugated capture antibodies with affinity to the target analyte. The assay format uses a standard ELISA sandwich with horseradish peroxidase-conjugated detector antibodies used to turnover a fluorogenic enzyme substrate in the reaction volume and accumulate fluorescent products (fig. 1a). The compartments are formed spontaneously through the addition of oil to the amphiphilic particles (Sahin et al., 2022; Wu et al., 2020). The inner PEG region stabilized a nanolitre droplet in which the enzyme-catalysed fluorescent product of the ADHP substrate is quickly concentrated to detectable levels.

**Figure 1:**
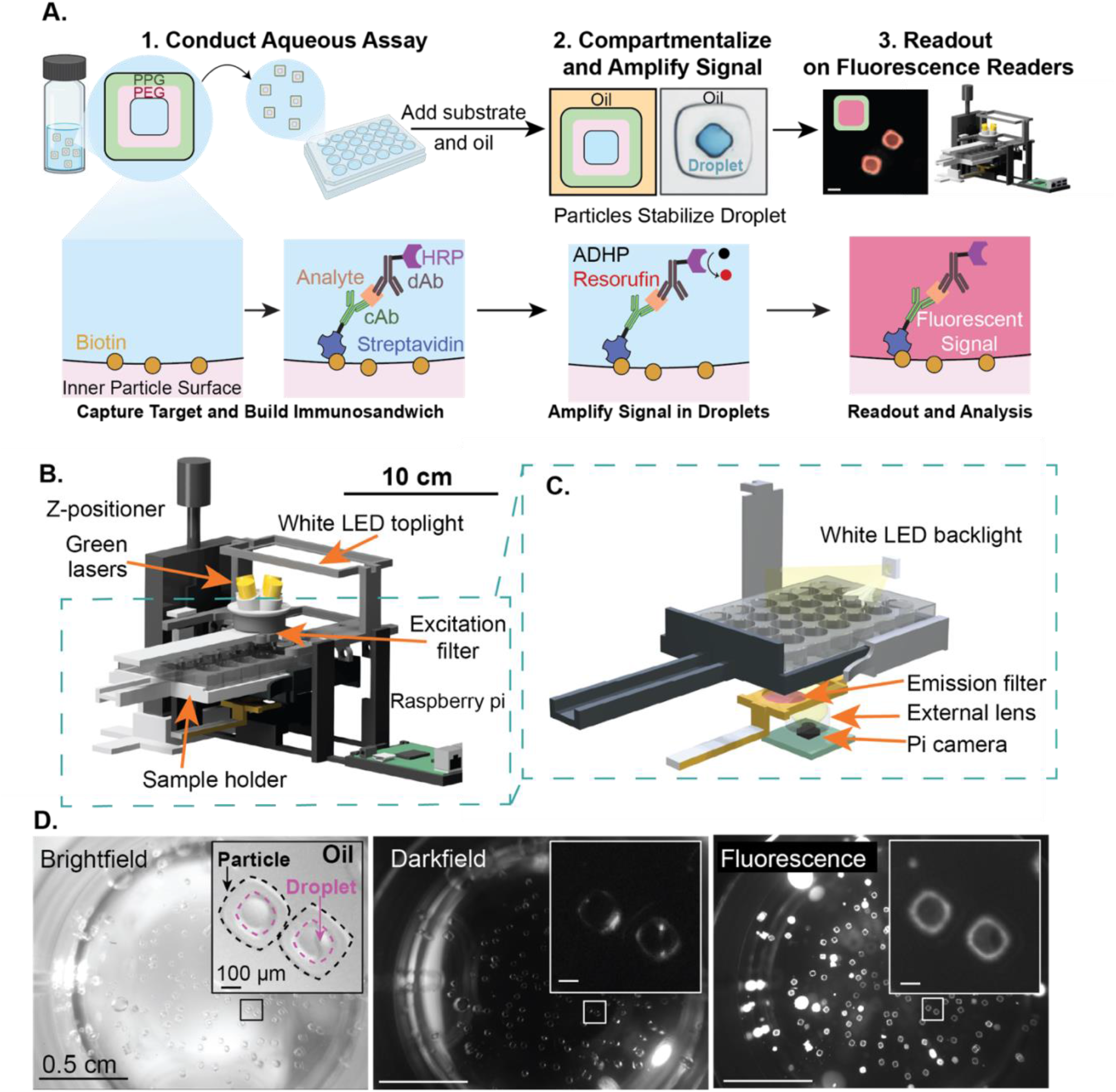
Amphiphilic Particles Workflow and Portable Reader Design. **(A)** Schematic of the lab on a particle workflow for performing and reading out parallel amplified immunoassays in droplets. Initial steps to form an immunosandwich complex are similar to traditional particle/bead based immunoassay platforms. After the final wash step, enzyme substrate and oil are added in quick sequence to isolate particle-templated droplets as individual reaction compartments. Fluorescence signal from enzymatic turnover of the ADHP substrate to resorufin quickly accumulates to detectable levels inside the small droplet volumes. The readout of each compartment is then conducted on a multimodal portable microscopic reader. **(B)** The multimodal reader design showing a Raspberry pi controlled automated camera for image acquisition, external lens capable of capturing the entire well field of view, excitation and emission filters for fluorescence acquisition, and LED backlights used for brightfield and darkfield illumination. **(C)** Magnified view of the imaging components of the reader showing laser diodes, emission filter, lens, and camera used to acquire images. **(D)** Representative brightfield, darkfield, and fluorescence images showing the whole well filled with particles following an assay (insert).

### Portable Reader and Image Processing Workflow

We designed a portable, low-cost, multi-modal reader to facilitate the readout from the assay (fig. 1b). The reader has a small footprint (10” x 8”) and consists of a z-positioning stage to focus sample, sample holder, Raspberry Pi-based controller, and an imaging module. The imaging module comprises 4 green (532 nm) laser diodes to excite fluorophores (570 nm peak), top white light emitting diodes (LED) for brightfield imaging, and a side angle LED for darkfield imaging (fig.1c). The images are captured with and without an insertable emission filter (used for fluorescence imaging), an external lens, and a Raspberry Pi camera. The magnification of the optical system, ∼0.2x, was designed to take whole well images in brightfield, darkfield, and fluorescence modalities (fig. 1d, and Methods).

We also developed an image processing pipeline (fig. 2) to automate microparticle detection and fluorescence intensity measurement for each droplet from captured images. We first created a high dynamic range (HDR) image from four individual low dynamic range (LDR) images. We then corrected the chromatic aberrations on brightfield and darkfield images and registered them to form a synthesized HDR image. A customized particle detection algorithm was applied to registered images for each modality individually, creating three binary masks which were then fused to one final detection mask. We noticed that due to the variations in lighting conditions in the different images taken of the same well, the algorithm detected different fractions of particles with each imaging mode (supplementary fig. 2). On average, the algorithm detected 24% of the particles from the brightfield image, 21% from the darkfield image and 56% from the HDR fluorescence image. The detectable percentage for HDR images was lower when the sample’s fluorescence signal was lower and vice versa. Combining masks from each individual brightfield, darkfield, and fluorescence image led to a majority of particles being detected, providing more measurements per sample. Each unconnected region of interest (ROI) was considered a single particle, and the fluorescence intensity profile was measured by overlaying the masks on the HDR image. Detecting from three modalities increased the number of particles that could be analysed, which can increase the accuracy and robustness of our system as described below, especially when fluorescence signals were low and contained insufficient information for detection. Due to a large number of particles initially seeded into a well (∼150), 100% particle detection accuracy is not needed for a successful assay readout.

**Figure 2:**
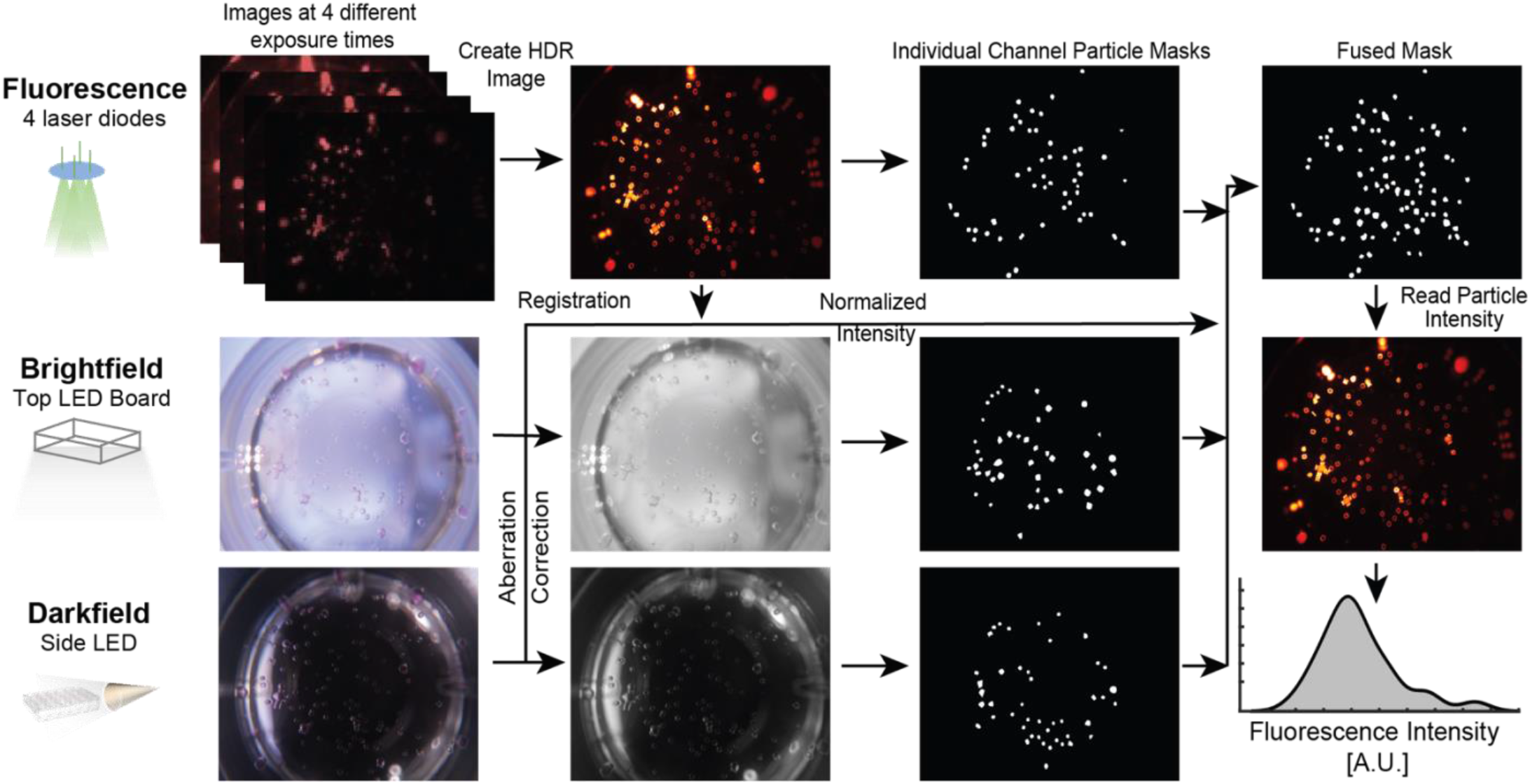
Image Processing Workflow. Fluorescence, brightfield, and darkfield images are captured following illumination with laser diodes, a top LED board, and side LEDs respectively. All acquired images are then fed into a customized image processing algorithm and fluorescence intensity measurements associated with each detected particle are returned. A pre-processing module first synthesizes the high-dynamic range (HDR) image, corrects the chromatic aberration, and registers images from first three modalities. Then, for each modality, an edge detector based algorithm extracts the regions of interest (ROI) containing particles and generates binary masks for the particles. Three binary particle masks are fused together through OR operations and the fluorescence intensities of each particle from the HDR image is assigned to each region of interest. The histogram shows the kernel density of fluorescence intensities from a population of particles.

### NT-proBNP Detection Using Amphiphilic Particles and Portable Reader

We first developed image analysis procedures for the combined assay and reader that maximized the capability to measure NT-proBNP levels above and below the clinically-relevant cut-off (∼ 0.1 ng/ml) concentration in buffer. We identified 3 potential regions of interest (ROIs) within a particle-droplet compartment to quantify fluorescence intensity (supplementary fig. 3): the inner PEG region, the droplet region, or the union of both the PEG and droplet regions (combined ROI). We found that the droplet region had the least fluorescence intensity out of the three ROIs and observed that resorufin, the fluorescent product of the enzyme-catalyzed oxidation of ADHP, appeared to accumulate or have higher fluorescence intensity in the PEG region. The PEG ROI and combined ROI had similar fluorescence intensities at 0.1 ng/ml (supplementary fig. 2), however using the combined ROI we were able to achieve higher statistical significance, p-value 10^−11^ versus 3×10^−9^, between the fluorescence intensity distributions of a negative control (0 ng/ml) and 0.1 ng/ml sample (supplementary fig. 3). As such, we used the combined ROI for all subsequent experimental analysis.

After establishing image analysis protocols, we characterized the assay performance by spiking various concentrations of NT-proBNP in buffer. Leveraging the ability to achieve multiple measurements per sample across an assembly of separate particles we could increase overall measurement accuracy, a process referred to as “swarm sensing” (Ouyang & Di Carlo, 2019). On average, we measured intensities from 80 particles per sample. Based on the distributions of fluorescence intensities of a negative control we set a threshold of mean + 3X of the standard deviation (*µ*_*negative control*_ + 3*σ*_*negative control*_) (fig. 3b inset). We then calculated the fraction of total particles detected with fluorescence intensities above this threshold for each NT-proBNP spiking concentration of 0.1, 1, and 10 ng/ml (fig. 3b). Across three repeats of this experiment, we found that the assay showed a step response, where the mean fraction above the threshold was close to zero (0.025) for 0 ng/ml, and was saturated at ∼1 for 0.1, 1, 10 ng/ml, accounting for variations across the experimental repeats (fig. 3c). The mean fluorescence intensity of the distributions across experimental repeats showed a dose-response type behavior with a linearly increasing mean fluorescence intensity with increasing NT-proBNP concentrations (fig. 3d). Overall, the assay was able to distinguish between the clinical cut-off at 0.1 ng/ml and negative control (0 ng/ml), with linearity in fluorescence intensities helping quantitate across a concentration range that is clinically meaningful for improved age-adjusted positive predive value (PPV) (0.45 – 1.8 ng/ml) for HF (Chow et al., 2017; Ezekowitz et al., 2017; Ponikowski et al., 2016).

**Figure 3:**
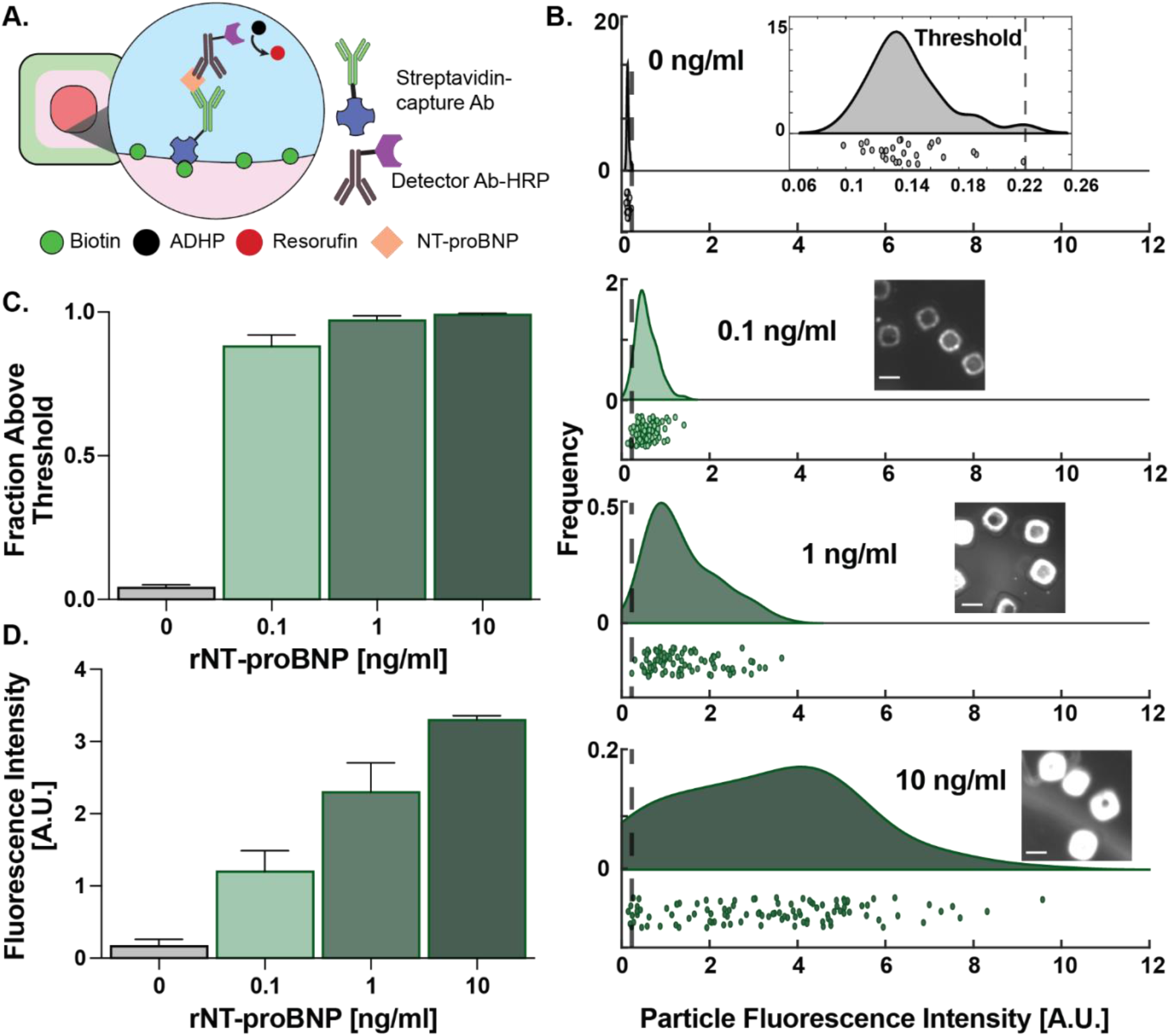
NT-proBNP Detection in Buffer. **(A)** Schematic showing immunosandwich formation for NT-proBNP detection on inner particle surface. **(B)** Raincloud plots showing distribution of fluorescence intensities of individual particles with increasing spiked NT-proBNP levels. Top insert shows threshold determination as µ_0_ + 3σ_0_ other inserts showing images of fluorescence signals in particle-droplets at respective concentrations. Scale bar is 100 μm. **(C)** Mean fraction of particles above threshold. Error bars show standard deviation. **(D)** Mean particle intensities from three experimental repeats.

Next, we sought to assess assay performance in a complex medium. We spiked NT-proBNP in plasma-EDTA samples previously depleted of endogenous NT-proBNP (<15 pg/ml) and diluted spiked plasma in buffer at a ratio of 1:3 to reduce matrix effects. The spiked samples were tested using the lab on a particle assay and analyzed by the portable reader. We noticed a correlated increase in fluorescence intensity with increasing spiked NT-proBNP (fig. 4a,b; supplementary fig. 4), with a linear response between 0.2 – 2 ng/ml and limit of detection (LOD) at 50 pg/ml (fig. 4b). Each individual particle-droplet provided an individual measurement, providing ∼80 measurements per sample; this multiplicity in measurement helps overcome random errors and improves quantitation with a low-cost reader. When the number of particles measured is increased from 3 – 80, the standard error of the mean for each measurement decreases from ∼27% to <5% (fig 4c), lowering error 5 fold on average across conditions.

**Figure 4:**
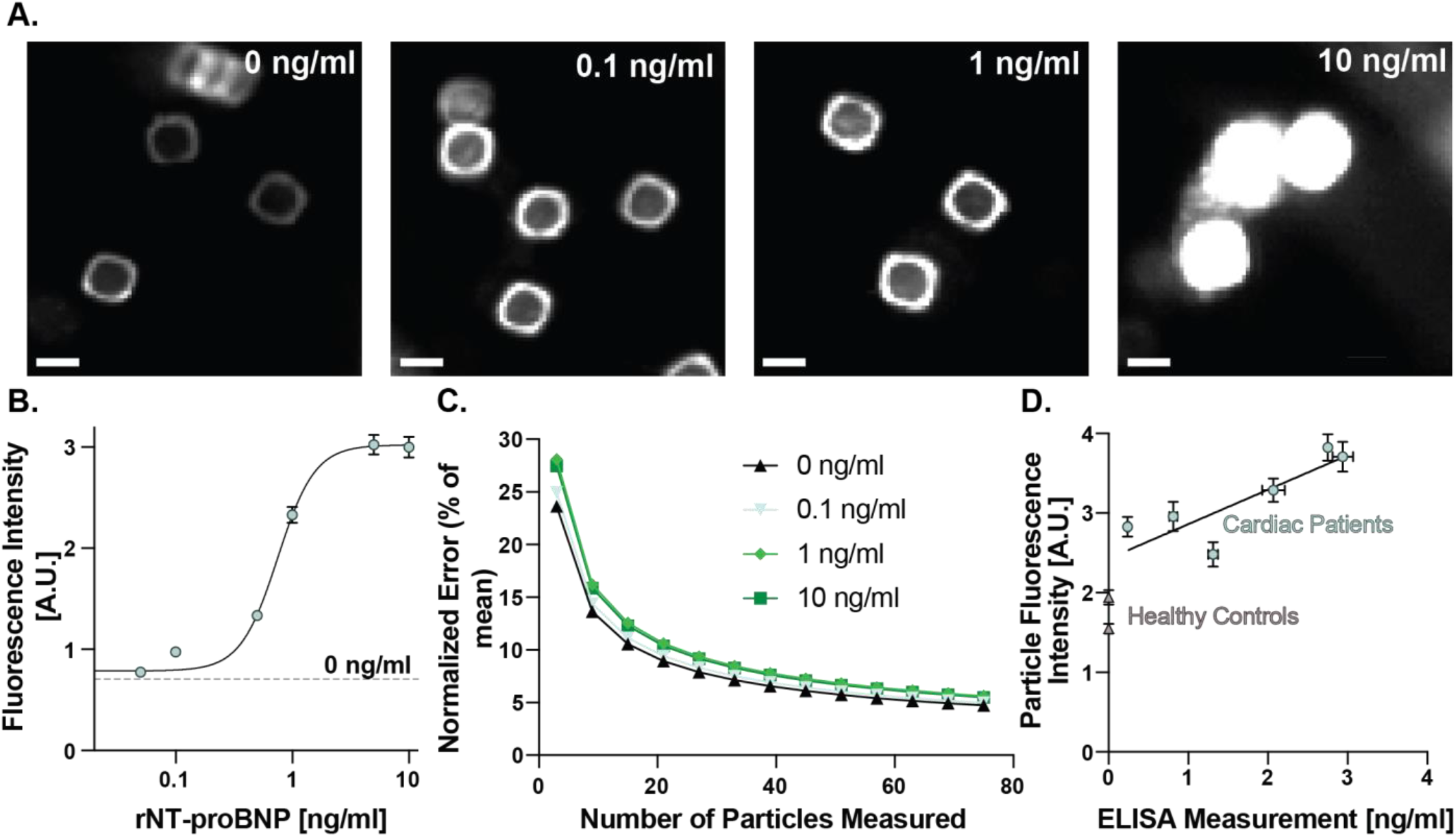
Plasma-EDTA spiked and patient testing. **(A)** HDR images showing concentration dependent response in fluorescence intensity. **(B)** Mean fluorescence intensities across 3 experimental repeats and >250 particle-droplets. **(C)** Standard error of the mean as a function of measurements taken per sample for varying conditions. **(D)** Measurements of cardiac patient samples using particle-portable reader system (y-axis) versus standard well plate based ELISA (x-axis). Control samples were below the quantitation threshold of the ELISA (<15 pg/ml).

To better understand the diagnostic impact of multiple measurements of the same sample by multiple amphiphilic particles, we conducted Monte Carlo simulations to understand the error rate of designating a sample as positive or negative based on counting the number of particles above a threshold set by the distribution of negative control particle intensities (supplementary fig. 5). In our analysis, we found that based on this thresholding method, the negative control sample had no particles above the threshold and was always designated as a negative. However, the sample at the clinical cut-off (0.1 ng/ml), showed a large variation (supplementary fig. 5a) based on the number of particles analyzed for that sample. As the number of measurements per sample increased, the error rate of designating a sample as false negative exponentially decreased, i.e. the ability of this low-cost system to correctly identify a sample at the cut-off as being positive asymptotically increased with the increasing number of particles measured. Across three experimental repeats (supplementary fig. 5b), we found that this thresholding method could easily differentiate between samples at or above the clinical cut-off (0.1 – 10 ng/ml) and those below (0, 0.05 ng/ml). This ability to obtain tens of measurements per sample and generate thresholds or summary statistics is a unique advantage of our particle-based assay system, which helps improve quantitative capabilities with a low-cost reader.

Following the serum characterization, we performed testing on a small pool of patient samples for which we had gold standard NT-proBNP levels by standard plate ELISA (supplementary fig. 6). The patient samples were well characterized by a gold standard ELISA measurement showing a low variation over 2 and 4 fold dilution curves (supplementary fig. 6a), with NT-proBNP concentrations ranging from 0.236 – 2.9 ng/ml. The particle and portable reader system results were well correlated with those of the ELISA measurement (fig. 4d), showing a linear correlation between the two measurements. Again, owing to the “swarm” sensing mechanism, the particle and portable reader system was able to achieve lower variations in measurements across the six patient samples (∼5%) compared to the standard ELISA for the same patient samples (∼9%) (supplementary fig. 5d) readout on a standard plate reader.

## Conclusion

Compartmentalized assays leverage small volumes to achieve higher sensitivity, however, they require significant infrastructure. Our amphiphilic particles can thermodynamically stabilize nanoliter droplets as reaction compartments foregoing the need for expensive equipment to form droplets or fill microchambers, and realizing compartmentalized assays in an accessible well plate format. Here, we combined this technology with a low-cost portable reader and image analysis algorithm to potentially extend the access of this technology for distributed use in local labs and clinics. We demonstrated the ability of this combined assay-reader system to measure NT-proBNP levels, a canonical heart failure marker, in complex media and patient samples. From the plurality of particles added per well, we were able to obtain tens of measurements per sample, thereby improving quantitation capabilities and reducing measurement error from using a low-cost imaging system. Particle numbers can also be adjusted depending on the application, and sensitivity requirements. In the future, we can also combine this with different shapes of particles to barcode each patient or biomarker and enable patient pooling or multiplexed analyte panel testing, for scaling and further lowering cost per patient per analyte. Computer vision algorithms can also help detect these barcodes from a single white light image, enabling barcoded droplet assays in this easily deployed format. In conclusion, this platform potentiates nanoliter droplet assay for screening and monitoring, thereby enabling earlier detection and quicker treatment/intervention decision times in low-resource settings.

## Methods

### Particle Fabrication and Functionalization

Coaxial flow lithography was implemented to fabricate amphiphilic particles using an improved 3D printed device design that is 3.5x smaller by volume than the previously reported devices, possible by advances in micro stereolithography (Boston Microfabrication). The new design also features angled, rather than perpendicular connections between inlets and internal channels, which along with the smaller channel dimensions decreases the chance of air bubbles being entrapped which would previously often lead to flow profile deformations (supplementary fig. 1). As previously reported (Destgeer et al., 2021), we flowed four precursor solutions through the concentric channels to obtain co-axial flows as follows from outer to inner stream: (1) inert sheath of poly(propylene glycol) [PPG], (2) poly(propylene glycol) diacrylate [PPGDA] + photo initiator, (3) poly(ethylene glycol) diacrylate [PEGDA] + Acrylate-PEG-Biotin + photo initiator, and (4) inert sheath of poly(ethylene glycol) [PEG]. Once the co-axial flow is created, we polymerized the reactive streams using UV light exposed through patterned slits in a photomask to obtain particles with defined 3D geometries. We can fabricate 100-120 particles per cycle, depending on the exposed region size. After fabrication, the remaining precursor material is washed away from particles by dilution using ethanol. Particles are stored in ethanol at 4°C before use.

### Amphiphilic Particle Assay

We first transferred biotinylated particles suspended in ethanol to a 24-well plate and counted them to ensure a similar distribution per well. Following the transfer, we washed the particles with PBSP (PBS with 0.5% w/v Pluronic F-127) before incubating with 10µg ml^-1^ streptavidin (ThermoFisher Scientific) for 30 minutes. After incubation, we washed the particles with PBSP and incubated with 10µg ml^-1^ capture antibody (15C4cc, HyTest) solution. Following another round of washing, we blocked the particle and plate surface using a protein-free blocking buffer (ThermoFisher Scientific) for 1 hour.

We incubated functionalized particles with samples, NT-proBNP (HyTest) at desired concentrations spiked in buffer or NT-proBNP free plasma-EDTA (HyTest). Control samples were incubated with only buffer or NT-proBNP free plasma. Following incubation with the antigen, we washed particles with a 0.05% Tween 20 solution to remove unbound antigens. We then incubated for an hour with 1.5µg ml^-1^ of HRP conjugated detection antibody (13G12cc, HyTest) in BSA buffer (PBS with 0.1% w/v BSA) and washed. For the signal generation, we added a QuantaRed assay solution (ThermoFisher) to the particles, immediately removed, and added PSDS oil (Sigma-Aldrich) to form aqueous droplets stabilized within the particles. The particles were incubated for signal development before imaging using the portable reader. We captured brightfield, darkfield and four fluorescence images (at 100, 500, 1000, and 2000 ms exposure) of each well.

### Portable Imager

The customized reader consisted of three major compartments: an illumination module, an imaging module, and a sample holder module. The illumination module had three independent illumination sources, top white LED board (Adafruit White LED Module, ID1622), four laser diodes (Q-BAIHE, Industrial Green Laser, 532MD-100-HS-GD) with a diffuser, and a white LED (SunLED XSFWCB983W-ND). The white LED board provided uniform white illumination for brightfield imaging. The side-diffused LED was placed so that the sensor was close but perpendicular to its light path. With the presence of the sample, some part of the illumination was scattered by the particles into the sensor, emphasizing the high-frequency signal (i.e., the PEG layer of the particles) while reducing the background. The four laser diodes were installed 15 degrees off the vertical axis and the focusing lenses were removed to form four uniform light cones, creating four overlaid elliptical light spots for a uniform illumination covering the entire well. The imaging module included a Raspberry Pi camera module with an embedded Sony IMX219 sensor, an external lens, and an insertable band-pass filter (Edmund Optics 625nm CWL, OD 4.0 25nm Bandpass Filter) to filter out the excitation light for the fluorescent channel. The sample holder module consisted of a z-positional for focusing and a slidable sample holder. Illumination sources and image acquisition were fully powered and controlled by a Raspberry Pi with a customized Graphical User Interface (GUI), except for the laser diodes which were powered by an external source. We applied heat paste around the laser diodes to avoid laser overheating. All holders for illumination and the sample well plate were customized and 3D-printed (Objet30 Pro, Stratasys Ltd.) The fluorescence images were read directly from the sensor in raw format. The raw format files are decoded and demosaiced to extract only the red channel. This process removed the redundant in-device pre-processing and reduced the laser operating time by ∼3-fold. Using raw images for fluorescence channels also improved the dynamic range of the sensor from 8-bit depth to 10-bit depth. Note that 8-bit fluorescence images suffer from several under- and over-exposure problems, and for 10-bit images the information loss was less significant but still occurred at negative control and 10 ng/mL concentrations.

### Image Acquisition

For each experiment, we first previewed the field-of-view using the brightfield channel and manually focused the images with the z-stage. We then captured fluorescence, brightfield, and darkfield images using four green laser diodes with a diffuser, top board white LED, and side LED as light sources, respectively. Four fluorescence images of each field-of-view were acquired with exposure times of 100, 500, 1000, and 2000 ms, where 2000 ms was the maximum exposure time of the sensor. All four fluorescence images were registered and used to generate the HDR image. The exposure time for brightfield and darkfield images was 50 ms. All image sizes are 3280 × 2464 pixels.

### Image Processing

Fluorescence images with different exposure times were registered using phase correlation (Reddy & Chatterji, 1996) to correct possible shifts among the images. A Gaussian-weighted HDR algorithm was then applied to synthesize the HDR image from LDR images.(Robertson et al., 1999) The brightfield and darkfield images were converted to greyscale, and corrected to minimize the effects of chromatic aberration with empirical second-order parameters. Then the corrected images were registered to the HDR image using the rigid phase correlation method (Srinivasa Reddy & Chatterji, 1996). We then preprocessed the brightfield and darkfield images using adaptive histogram equalization (Zuiderveld, 1994) and bilateral Gaussian filtering (Tomasi & Manduchi, 1998) to enhance the contrast and reduce noise while preserving the edges. To detect particles from images, we utilized a Canny edge detector to extract the particle boundaries. (Canny, 1986) Image dilation was then applied to binary images from the Canny detector, and morphological processing was utilized to fill the holes in the images and generate masks for particles. Each unconnected mask represented a potential detected region of interest (ROI) for a particle but included false positive detections. False positive masks were filtered out based on their area (size) and eccentricity (shape) with empirical thresholds. Overlapped particles and particles near the edge of the well were excluded. This detection process was fine-tuned and applied to each modality individually. Then the three binary masks were fused together with a convexity constraint. The fused mask was superimposed on an HDR image to define ROI, and fluorescence intensities from these ROIs were measured.

### ELISA testing

To determine the gold standard NT-proBNP concentrations, we performed ELISA on patient samples using a kit (Abcam ab263877). The kit included a 96-well plate with capture antibodies pre-bound to each well. Patient serum samples were analyzed whole, half and quarter diluted. After incubation with both patient samples and then detection antibodies, the colorimetric optical density readout was done at 450nm on a microplate reader.

### Monte Carlo Swarm Sensing Simulations

To study the effects of number of particles detected on NT-proBNP quantification, we applied Monte Carlo simulation to evaluate the variation and robustness of the measurements. We first randomly selected an experiment from the serum sample, and extracted the intensities profiles of all detected particles for negative control (0 pg/ml) and clinical cut-off (100 pg/ml). Then we randomly sampled a subset of all intensities (from 1 to 50 particles) for both concentrations. For the selected intensities, we measured the mean intensity for each concentration and calculated p-value for a single-tailed t-test and fraction above threshold. The random sampling was repeated 5000 times with uniform probability for each particle, and we measured the mean and standard deviation for all three metrics across the repeats.

## Aknowledgement

The authors would like to acknowledge the National Science Foundation PATHS-UP Engineering Research Center (Grant #1648451)

