## Supplementary Figures for "Amphiphilic particle-stabilized nanoliter droplet reactors with a multi-modal portable reader for distributive biomarker quantification"


Supplementary Figure 1: Improved 3D printed co-axial flow fabrication device

Supplementary Figure 2: Particle Recognition on the Multimodal Reader

Supplementary Figure 3: Regional Intensity and ROI

Supplementary Figure 4: rNT-proBNP spiked in plasma-EDTA

Supplementary Figure 5: Reduction in measurement error due to multiple measurements per sample

Supplementary Figure 6: Patient Sample Validation

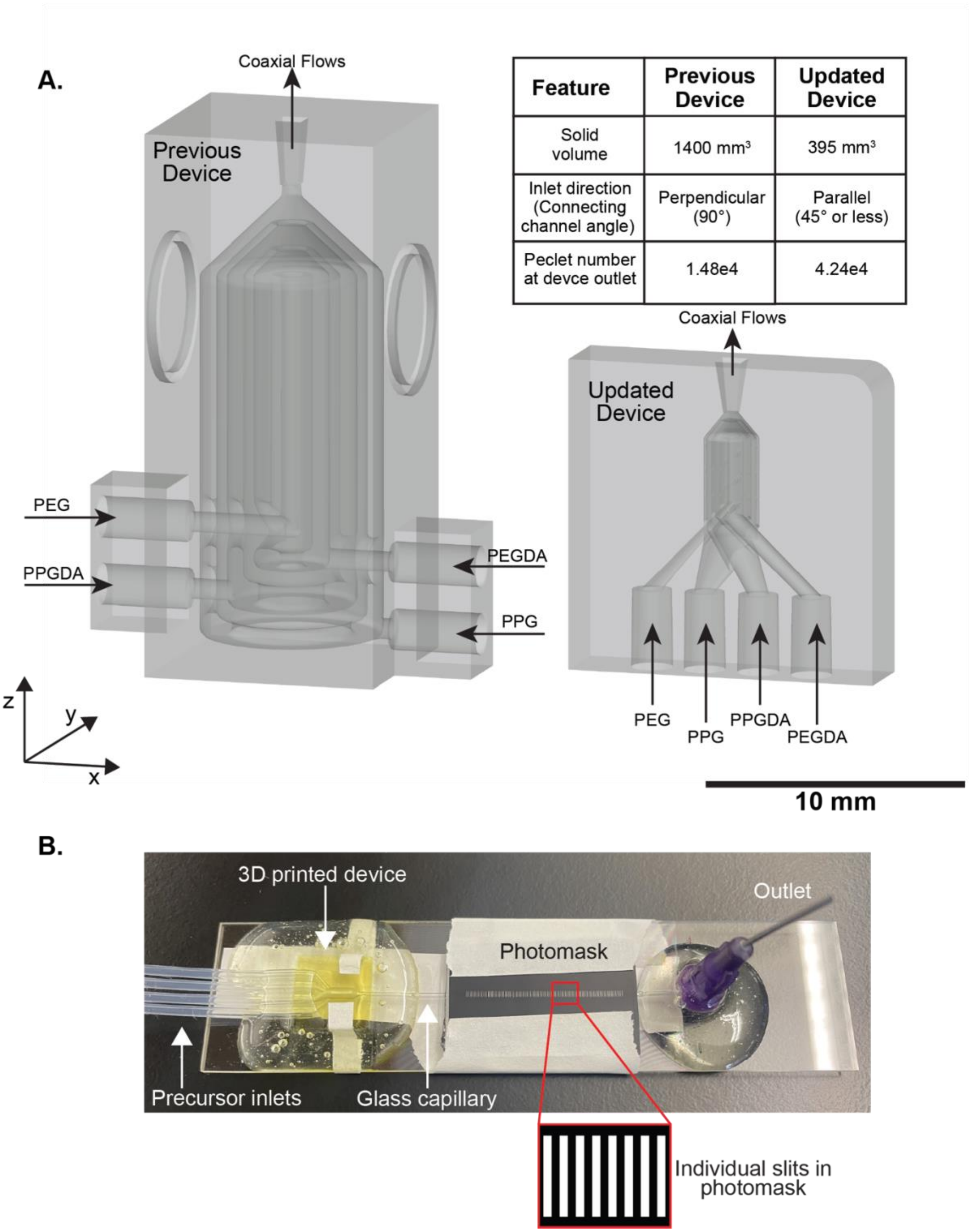


| Feature | Previous Device | Updated Device |
|---|---|---|
| Solid volume | 1400 mm³ | 395 mm³ |
| Inlet direction (Connecting channel angle) | Perpendicular (90°) | Parallel (45° or less) |
| Peclet number at devce outlet | 1.48e4 | 4.24e4 |

**Supplementary Figure 1: Improved 3D printed co-axial flow fabrication device. (A)** Comparison of previously published device (left) and improved, scaled down device (right) designs as CAD images. The new design is 3.5x smaller by volume and features inlets aligned with the direction of flow for more streamlined device assembly and fabrication. Peclet number is calculated at the point where outer two streams meet near the device outlet. **(B)** Image of an assembled device where the 3D printed device is connected to inlet tubing and polymerization capillary at the device outlet. A photomask is applied over the capillary, through which UV light can be applied.

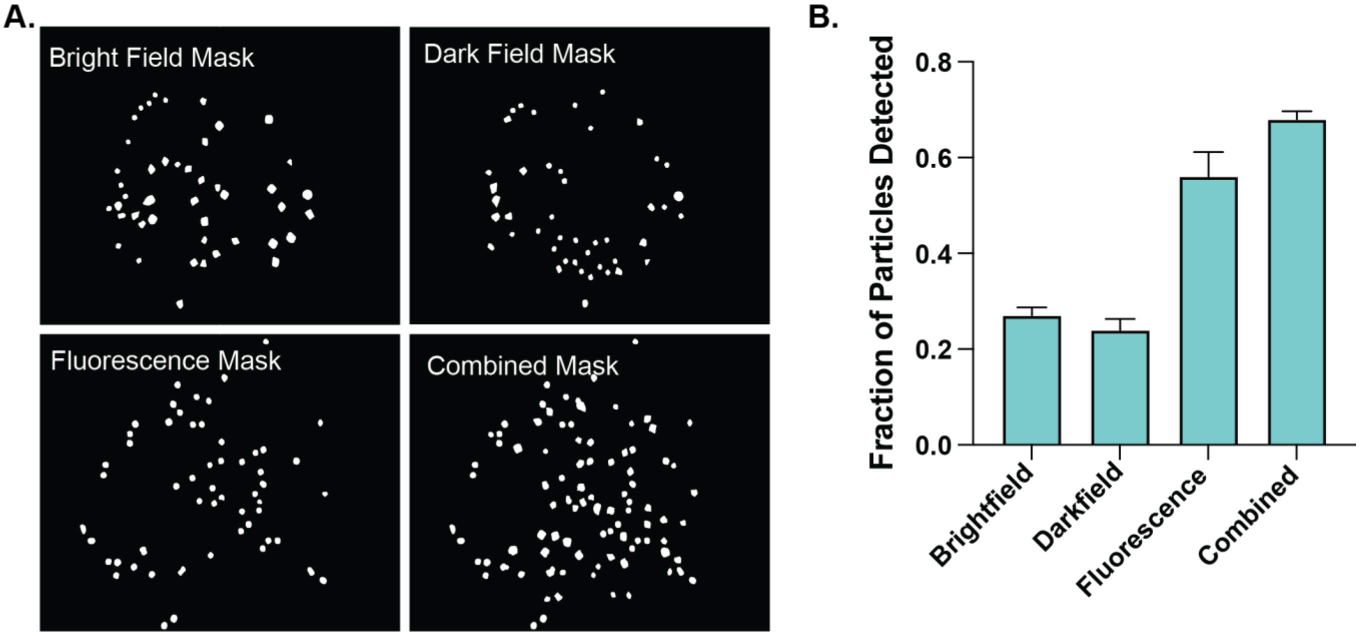


**Supplementary Figure 2: Particle Recognition on the Multimodal Reader. (A)** Images showing particle masks identified from individual brightfield, darkfield, and HDR fluorescence images, and a combined (fused) mask. **(B)** Combining particle masks from multiple imaging modalities enables detection of a larger fraction of particles per well. Data shown is from 3 repeats and error bars represent standard deviation.

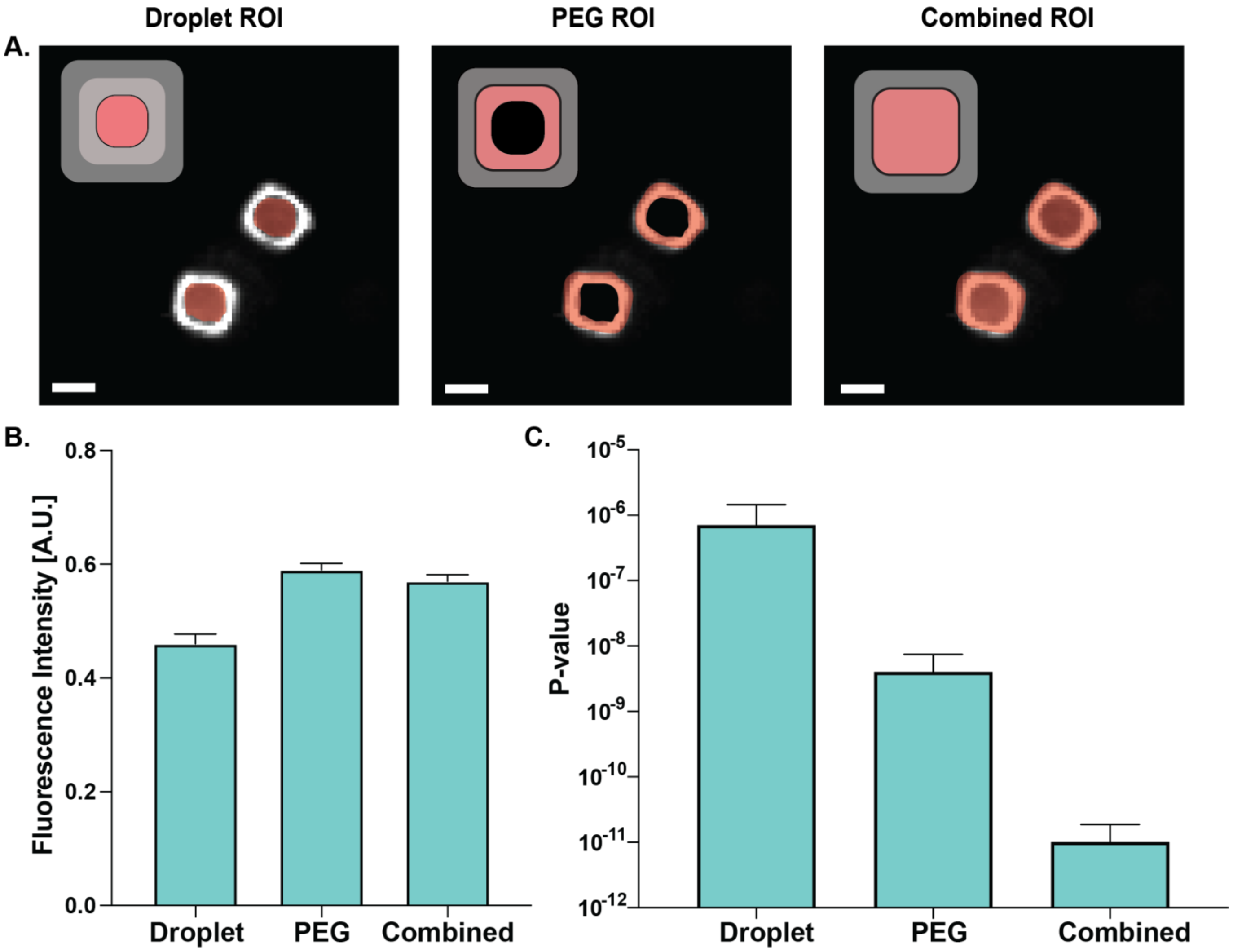


**Supplementary Figure 3: Regional Intensity and ROI. (A)** Three different types of ROI enabled by the lab on a particle assay are schematically shown. The droplet ROI encompasses only the internal droplet in the particle cavity, the PEG ROI consist of only the inner PEG region of the amphiphilic PPG/PEG particles, and the combined ROI includes both the droplet and PEG region in the fluorescence intensity measurements. **(B)** Mean intensity calculated from the three different ROIs for n = 82 particles at an NT-proBNP spike concentration of 0.1 ng/ml. Error bars show standard deviation. **(C)** Statistical significance between intensity distributions of negative control (0 ng/ml) and clinical cut-off (0.1 ng/ml) for each of the ROIs. Data shown is from 3 individual repeats, each repeat includes >50 individual particles detected. Error bars show standard deviation between repeats. The combined ROI enables the highest statistical significance in differentiating across the cut-off.

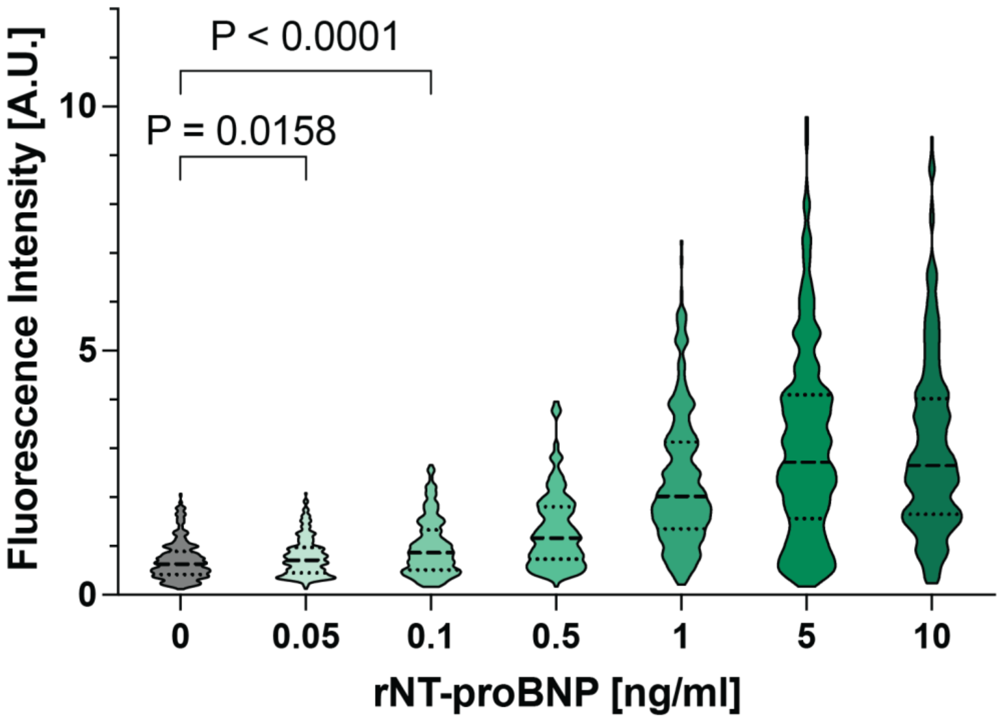


**Supplementary Figure 4: rNT-proBNP spiked in plasma-EDTA.** Violin plots showing distributions of particle intensities as a function of spiked rNT-proBNP concentration.

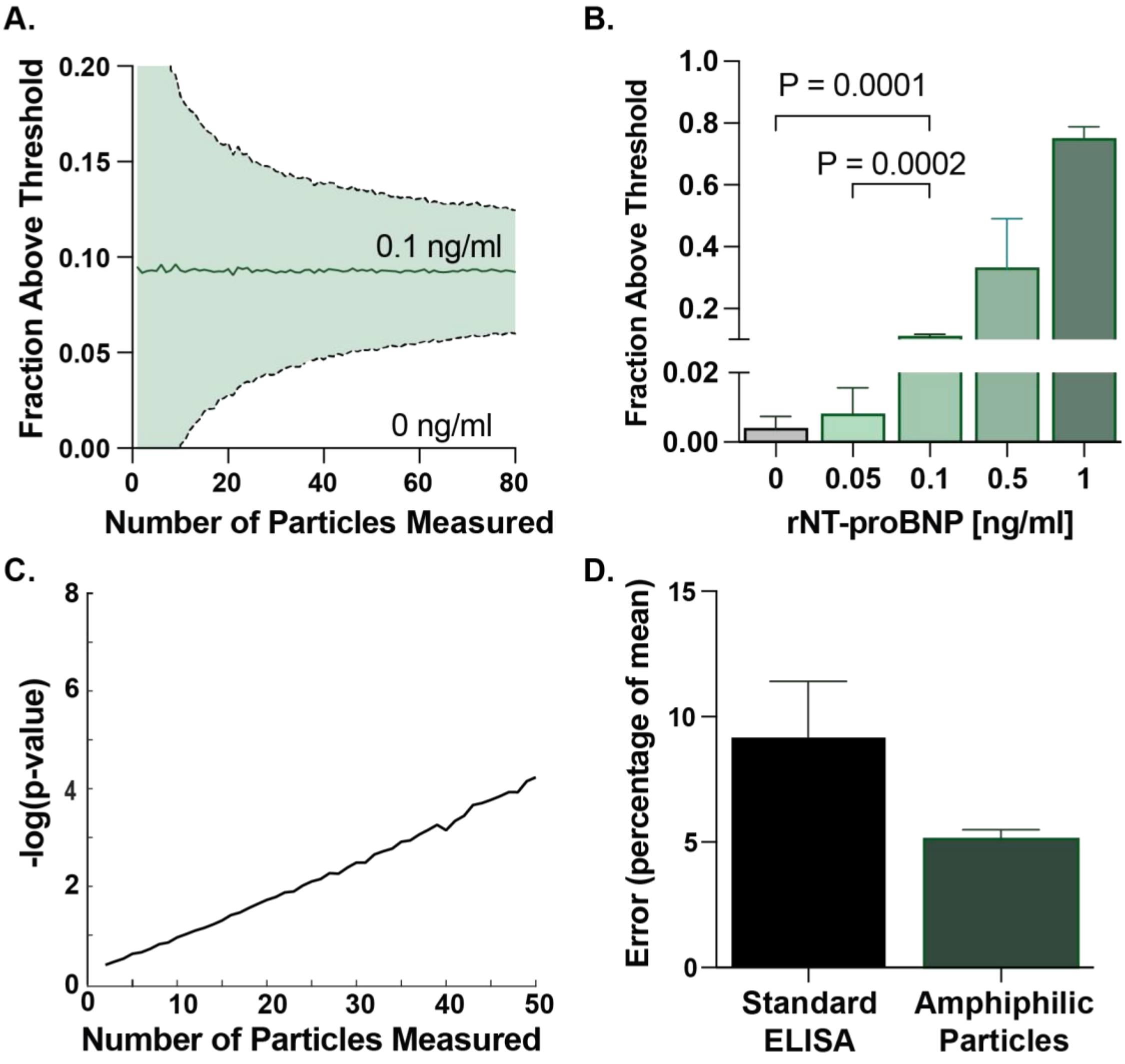


**Supplementary Figure 5: Reduction in measurement error due to multiple measurements per sample. (A)** Monte Carlo simulations (n = 5000) show how the variation in fraction of particles above threshold (set from distribution of negative control particle-drop intensities)**,** changes as a function of number of particles measured. As more particles are measured, the error rate of designating a sample at clinical cut-off (~0.1 ng/ml) as a false negative exponentially decrease. **(B)** Mean fraction above threshold for varying concentrations of NT-proBNP spiked in plasma-EDTA. Error bars show standard deviation. **(C)** Statistical capability to differentiate between distributions of 0 and 0.1 ng/ml increases as a function of particles measured per sample. **(D)** Mean variation (error) in measurement, from standard well plate based ELISA and amphiphilic particle based ELISA, across 6 cardiac patient samples

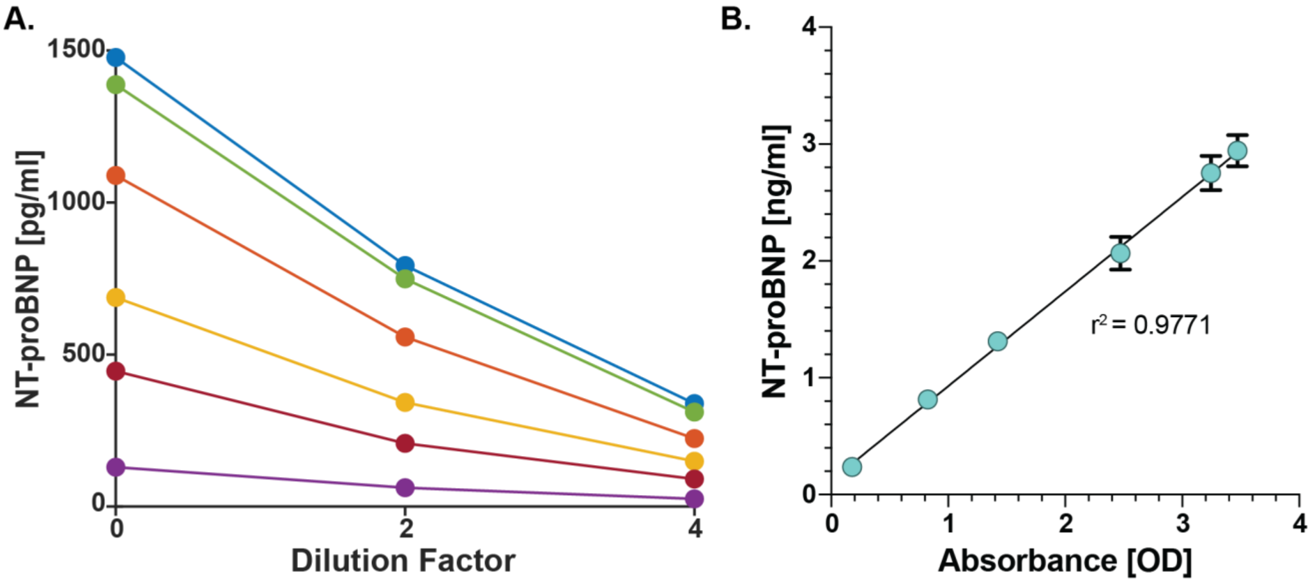


**Supplementary Figure 6: Patient Sample Validation. (A)** Dilution curves for the six patient samples with NT-proBNP levels > 100 pg/mL validated by external clinical ELISA chosen for patient testing on lab on a particle format. The two healthy samples had NT-proBNP levels below the limits of detection of the ELISA. **(B)** Mean and standard deviation of the ground truth measurements from patient samples within the linear range of the assay. These patient samples were chosen for testing on the lab on a particle platform since they had low variation in the dilution curves and produced linear response on the clinical validation test.
